# Inactivation of genes in oxidative respiration and iron acquisition pathways in pediatric clinical isolates of small-colony variant *Enterobactericeae*

**DOI:** 10.1101/662486

**Authors:** Alexander L. Greninger, Amin Addetia, Yue Tao, Amanda Adler, Xuan Qin

## Abstract

Isolation of bacterial small colony variants (SCVs) from clinical specimens is not uncommon and can fundamentally change the outcome of the associated infections. Bacterial SCVs often emerge with their normal colony phenotype (NCV) co-isolates in the same sample. The genetic and biochemical basis of SCV emergence *in vivo* is not well understood in Gram-negative bacteria. In this study, we interrogated the causal genetic lesions of SCV growth in three pairs of NCV and SCV co-isolates of *Escherichia coli, Citrobacter freundii*, and *Enterobacter hormaechei*. We confirmed the isogenic basis of SCV emergence, as there were only 4 single nucleotide variants in SCV for *E. coli*, 5 in *C. freundii*, and 8 in *E. hormaechei*, with respect to their NCV co-isolate. In addition, a 10.2kb chromosomal segment containing 11 genes was deleted in the *E. hormaechei* SCV isolate. Intriguingly, each SCV had at least one coding change in a gene associated with bacterial oxidative respiration and another involved iron capture. Chemical rescue confirmed the causal role of heme biosynthesis in *E. coli* and *C. freundii* and lipoic acid in *E. hormaechei* SCV isolates. Genetic rescue restored normal growth under aerobic conditions for *fes* and *hemL* in *C. freundii*; *hemL* in *E. coli*; and *lipA* in *E. hormaechei* SCV isolates. Prototrophic growth in all 3 SCV *Enterobacteriaceae* species was unaffected under anaerobic culture conditions *in vitro*, illustrating how SCVs may persist *in vivo* by abandoning the highly energetic lifestyle in an iron-limiting environment. We propose that the selective loss of functions in oxidative respiration and iron acquisition is indicative of bacterial virulence attenuation for niche specialization and persistence *in vivo*.

**Importance**

Small colony variant (SCV) bacteria are routinely isolated in the clinical microbiology laboratory and can be notoriously difficult to treat. Most studies of the genetic underpinnings of SCV clinical isolates have examined *Staphylococcus aureus* and few have looked at how SCV emerge in Gram-negative bacteria. Here, we undertook detailed characterization of three clinical isolates of SCV in *Escherichia coli, Citrobacter freundii*, and *Enterobacter hormaechei* along with their NCV co-isolates. Genomic sequencing revealed that each SCV had at least one coding change in genes involved in both bacterial oxidative respiration and iron capture. Chemical and genetic rescue revealed that both pathways could be responsible for the small colony variant. Each of the SCV showed no growth defect compared to NCV when incubated under anaerobic conditions, indicating a potential mechanism for SCV survival *in vivo*. We hypothesize that by retreating to anaerobic environments and avoiding escalating iron competition with the host, SCV have adapted to live to see another day.

## Introduction

To survive in the hostile host environment, bacteria may take two separate paths. The first and most commonly discussed is an arms race of iron competition and acquisition of antimicrobial resistance genes and pathogenicity factors (1–3). The alternative path is to adapt to persist through reductions in metabolic needs and an attenuated growth rate. The isolates that display this alternative phenotype are known as small colony variants (SCVs). Bacterial SCVs were first described in *Salmonella typhi* over a hundred years ago, prior to the antibiotic era (4). Isolation of SCVs is especially common in recurrent or persistent infections involving the respiratory tract, urinary tract, mid-ear, foreign body-related implants, and bone and joint (5–7). Emergence of bacterial SCVs from normal colony phenotype (NCV) parental isolates has been described in various clinical settings (4). Previously characterized bacterial SCV species include *Staphylococcus aureus, Escherichia coli, Neisseria gonorrhoeae, Stenotrophomonas maltophilia, Enterococcus*, and *Salmonella* (8–12). In addition to their decreased growth rate, bacterial SCV isolates are characterized by auxotrophism for components directly involved in the electron transfer chain, such as heme and menaquinone. SCVs may also display a nutritional dependency on thymidine or methionine (8, 13). From a treatment perspective, SCVs display a reduced response to antibiotics despite not carrying the associated antimicrobial resistance genes (4, 6, 14, 15). These degenerative changes allow SCV to persist *in vivo* in the unique environment and selection pressures of the infected host.

In the clinical microbiology laboratory, SCVs have been best characterized among *S. aureus* isolates. Common gene lesions in *S. aureus* SCVs have been seen in the heme, menaquinone, and thymidine biosynthetic pathways (6, 7). In clinical strains, disruptions in menaquinone biosynthesis have been associated with mutations in *menB, menC, menE* and *menF* (16). Laboratory-derived SCV *S. aureus, Salmonella typhimurium*, or *E. coli* with *hemA, hemB, hemD, hemL, lipA*, or *ctaA* mutations have been used for functional characterization of their changes in growth, metabolism, antimicrobial susceptibility, host invasion, and persistence (17–21). In addition, mutations in transcriptional regulators in *Staphylococcus spp*. governing bacterial virulence factor expression, such as *agr, sarA, sigB*, and *relA*, have also been detected, suggesting attenuated cytotoxicity may enable bacterial persistence (7, 15, 22, 23). However, the basis of SCV formation and persistence in clinical isolates of Gram-negative bacteria has been less well characterized (10, 24).

Laboratory recognition, isolation, characterization and appropriate report of bacterial SCVs have suffered from a lack of established standards or guidelines. We previously reported a method practical in clinical laboratories for recognition and phenotypic characterization of SCV *S. maltophilia* from airway secretions of CF patients (8). Our lab has since implemented a systematic, culture-based approach that checks not only for colony variation in size, texture, color, or hemolysis, but also inability to grow on the standard Mueller-Hinton (MH) medium for susceptibility testing. Using this systematic approach, we identified 3 pairs of clinical NCV and SCV *Enterobacteriaceae* co-isolates from blood and urine cultures. We then used whole genome sequencing to screen for the molecular mechanisms distinguishing the SCVs from their NCV co-isolates. Confirmatory chemical and genetic rescues were performed on the SCVs to determine which mutations were causal for the altered phenotype.

## Materials and Methods

### Isolation and characterization of SCV and NCV from clinical samples

This study was approved by Seattle Children’s Hospital Institutional Review Board. Three clinical NCV and SCV co-isolate pairs of *Escherichia coli, Citrobacter freundii*, and *Enterobacter hormaechei* were obtained from urine or blood cultures. Colony variants were separated by subcultures and bacterial species identification was performed using MALDI-TOF (Bruker biotyper, Bruker Daltonics, Inc.). Of note, the SCVs isolates described here did not grow on Mueller-Hinton agar and, thus, were reported in the patient’s clinical record.

### Case histories

*Case 1* - A previously healthy 6-week-old male with no history of hospitalization or receipt of antibiotics, presented to the emergency department with fever. His white blood cell count was elevated at 21,600/ml and a 2+ urine leukocyte esterase at the time of emergency visit. Given the patient’s age, the patient was admitted for a rule out sepsis workup and the patient was started on empiric ceftriaxone. Urine cultures grew 10^3^ - 10^4^ cfu/mL *Escherichia coli* and 10^3^ - 10^4^ cfu/mL SCV *Escherichia coli*. It was felt the urine culture did not support the diagnosis of UTI, and no additional antibiotics were indicated. The patient recovered fully.

*Case 2* - A 2-month-old female with right duplicated collection system presented to the emergency department with fever and foul smelling urine. The patient had experienced two urinary tract infections (UTIs) in the previous month (*Escherichia coli* and *Citrobacter* spp.) that were treated with amoxicillin and cephalexin, respectively. Given that the patient was currently receiving antibiotics for the previous Gram-negative UTI, the decision was made to admit the patient for likely IV antibiotic treatment, and the patient was started on empiric piperacillin-tazobactam. Her urine leukocyte esterase was 3+ with elevated red and white blood cells in urine at the time of culture. Urine cultures grew >10^5^ cfu/mL *Citrobacter* spp. and 5•10^4^ – 10^5^ cfu/mL SCV *Citrobacter* spp. and therapy was switched to ciprofloxacin. The patient completed 14 days of therapy and recovered fully.

*Case 3* - A 6 year-old male with end stage renal disease managed with renal dialysis presented to dialysis clinic with fever, hypotension, and tachycardia. The patient had a history of multiple bloodstream infections treated with ceftazidime, rifampin, gentamicin, or intravenous trimethoprim-sulfamethoxazole. The patient was admitted to the hospital and started on empiric vancomycin and gentamicin. Blood cultures grew out *Enterobacter cloacae* and SCV *Enterobacter cloacae* by MALDI-TOF and therapy was switched to cefepime. Additionally, thepatient received ceftazidime line-lock therapy. The patient completed treatment and recovered fully. Though the isolates were resulted out as *E. cloacae*, they were later identified as *E. hormaechei* based on whole genome sequencing.

### Whole genome sequencing of isolates

DNA was extracted from using the MoBio UltraClean Microbial DNA Isolation kit. DNA was diluted to 1ng/uL and tagmented using quarter-volume reactions of Nextera XT with 15 cycles of PCR amplification. Libraries were sequenced on a 2×300bp run on an Illumina MiSeq to achieve >1 million paired-end reads per sample. Reads were quality and adapter-trimmed using cutadapt (Q30) and de novo assembled using SPAdes using default parameters. Isolate details are reported in Table 1. For each pair of isolates, the NCV assembly was annotated using prokka and the reads from the paired isolate were mapped to the annotated assembly in Geneious v9.1. Variants were called using a minimum coverage of 7X and a minimum allele frequency of 75% (25). All variants were manually curated and variants within 10 bp of the edge of a contig were removed. All variants are displayed in Table 2 and all assemblies are available in NCBI BioProject PRJNA523376.

**Table 1.**
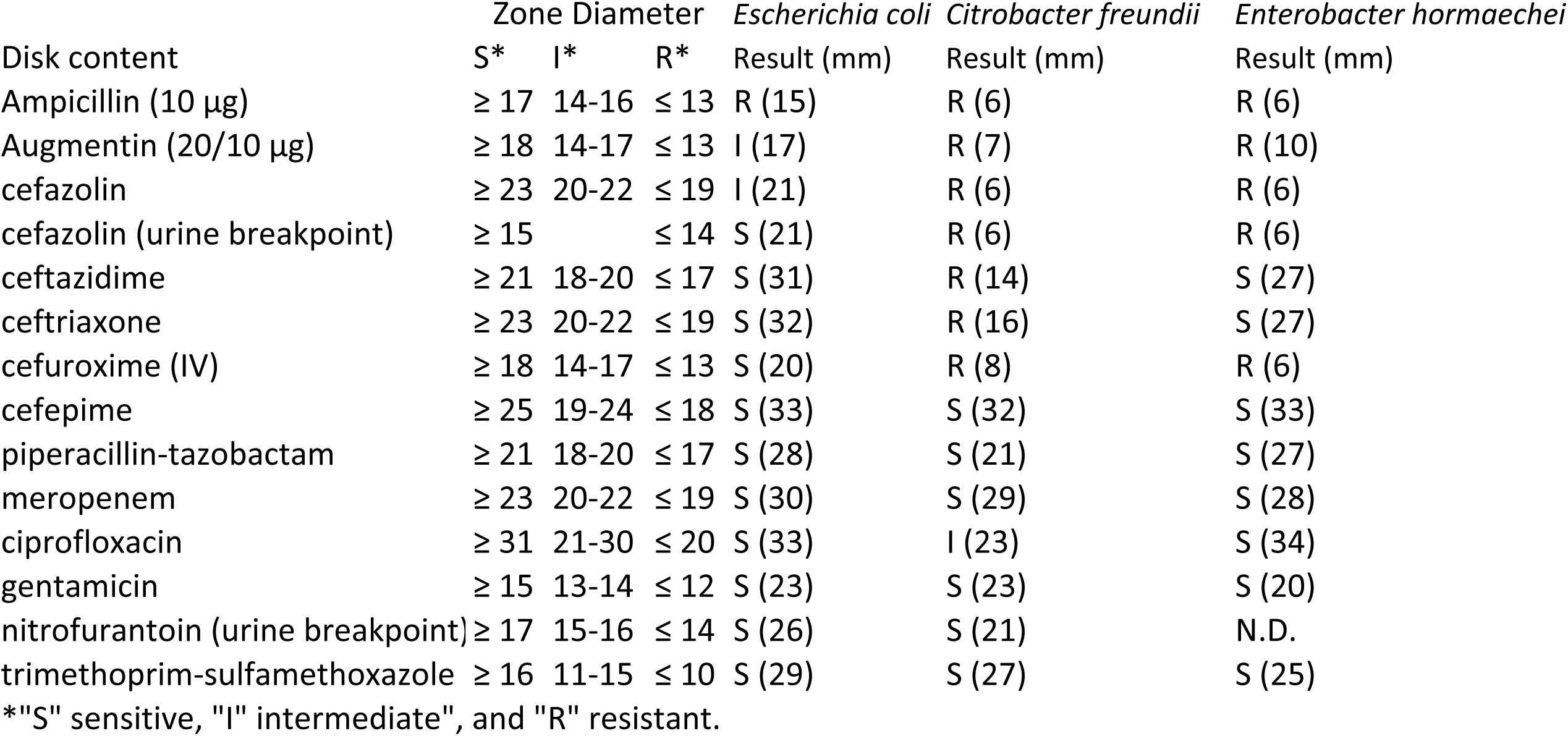
- Antibiotic susceptibility by disk diffusion of normal colony variant for each isolate based on CLSI M100-S29 breakpoints.

| Disk content | Zone Diameter |  |  | *Escherichia coli* | *Citrobacter freundii* | *Enterobacter hormaechei* |
| --- | --- | --- | --- | --- | --- | --- |
|  | S* | I* | R* | Result (mm) | Result (mm) | Result (mm) |
| Ampicillin (10 µg) | ≥ 17 | 14-16 | ≤ 13 | R (15) | R (6) | R (6) |
| Augmentin (20/10 µg) | ≥ 18 | 14-17 | ≤ 13 | I (17) | R (7) | R (10) |
| cefazolin | ≥ 23 | 20-22 | ≤ 19 | I (21) | R (6) | R (6) |
| cefazolin (urine breakpoint) | ≥ 15 |  | ≤ 14 | S (21) | R (6) | R (6) |
| ceftazidime | ≥ 21 | 18-20 | ≤ 17 | S (31) | R (14) | S (27) |
| ceftriaxone | ≥ 23 | 20-22 | ≤ 19 | S (32) | R (16) | S (27) |
| cefuroxime (IV) | ≥ 18 | 14-17 | ≤ 13 | S (20) | R (8) | R (6) |
| cefepime | ≥ 25 | 19-24 | ≤ 18 | S (33) | S (32) | S (33) |
| piperacillin-tazobactam | ≥ 21 | 18-20 | ≤ 17 | S (28) | S (21) | S (27) |
| meropenem | ≥ 23 | 20-22 | ≤ 19 | S (30) | S (29) | S (28) |
| ciprofloxacin | ≥ 31 | 21-30 | ≤ 20 | S (33) | I (23) | S (34) |
| gentamicin | ≥ 15 | 13-14 | ≤ 12 | S (23) | S (23) | S (20) |
| nitrofurantoin (urine breakpoint) | ≥ 17 | 15-16 | ≤ 14 | S (26) | S (21) | N.D. |
| trimethoprim-sulfamethoxazole | ≥ 16 | 11-15 | ≤ 10 | S (29) | S (27) | S (25) |

\*"S" sensitive, "I" intermediate", and "R" resistant.

**Table 2.**
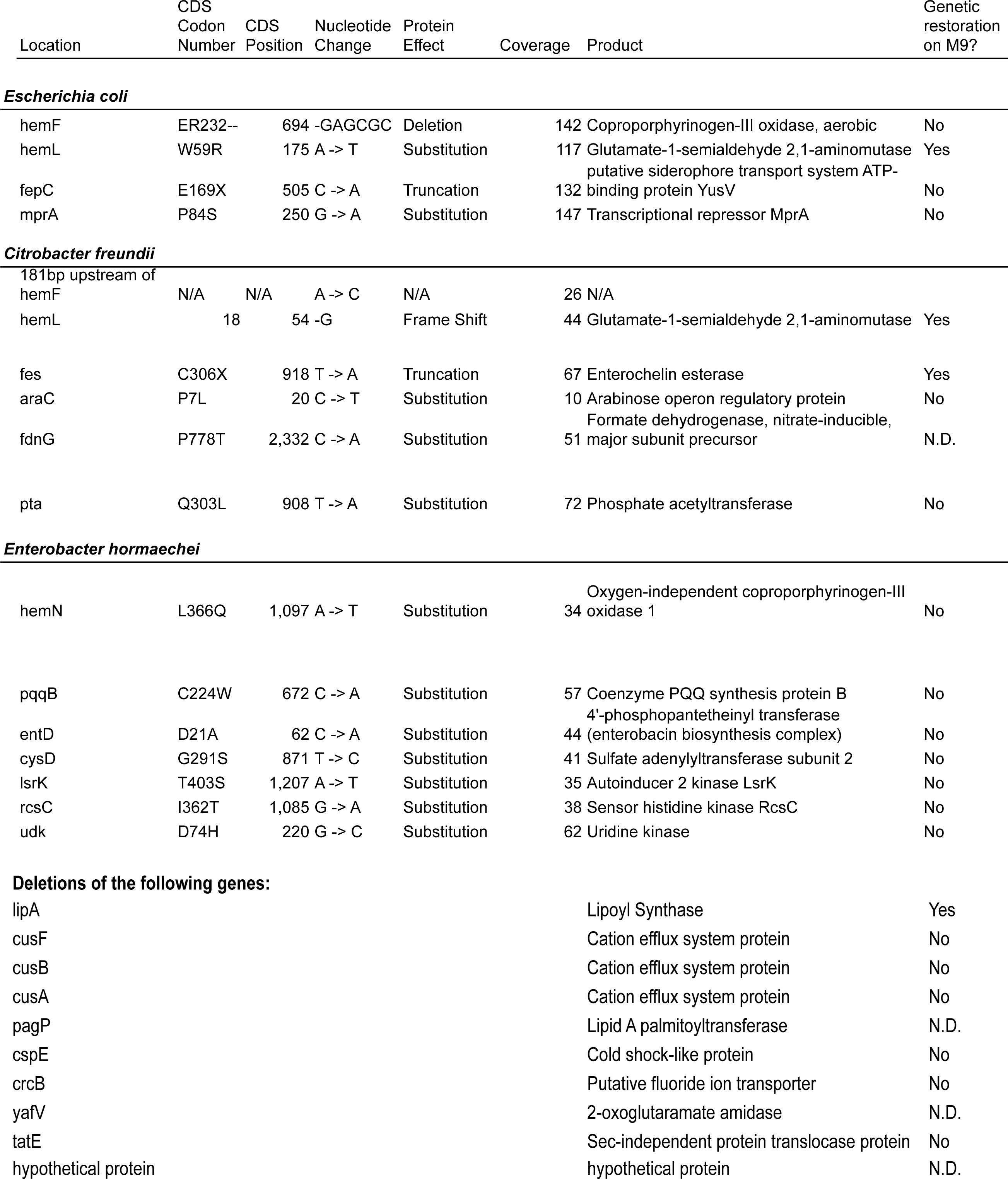
Genomic variants isolated from paired clinical co-isolates of Escherichia coli, Citrobacter freundii, Enterobacter hormaechei.

| Location | CDS<br>Codon<br>Number | CDS<br>Position | Nucleotide<br>Change | Protein<br>Effect | Coverage | Product | Genetic<br>restoration<br>on M9? |
| --- | --- | --- | --- | --- | --- | --- | --- |
| **Escherichia coli** |  |  |  |  |  |  |  |
| hemF | ER232-- | 694 | -GAGCGC | Deletion | 142 | Coproporphyrinogen-III oxidase, aerobic | No |
| hemL | W59R | 175 | A -> T | Substitution | 117 | Glutamate-1-semialdehyde 2,1-aminomutase | Yes |
| fepC | E169X | 505 | C -> A | Truncation | 132 | putative siderophore transport system ATP-binding protein YusV | No |
| mprA | P84S | 250 | G -> A | Substitution | 147 | Transcriptional repressor MprA | No |
| **Citrobacter freundii** |  |  |  |  |  |  |  |
| 181bp upstream of hemF | N/A | N/A | A -> C | N/A | 26 | N/A |  |
| hemL | 18 | 54 | -G | Frame Shift | 44 | Glutamate-1-semialdehyde 2,1-aminomutase | Yes |
| fes | C306X | 918 | T -> A | Truncation | 67 | Enterochelin esterase | Yes |
| araC | P7L | 20 | C -> T | Substitution | 10 | Arabinose operon regulatory protein | No |
| fdnG | P778T | 2,332 | C -> A | Substitution | 51 | Formate dehydrogenase, nitrate-inducible, major subunit precursor | N.D. |
| pta | Q303L | 908 | T -> A | Substitution | 72 | Phosphate acetyltransferase | No |
| **Enterobacter hormaechei** |  |  |  |  |  |  |  |
| hemN | L366Q | 1,097 | A -> T | Substitution | 34 | Oxygen-independent coproporphyrinogen-III oxidase 1 | No |
| pqqB | C224W | 672 | C -> A | Substitution | 57 | Coenzyme PQQ synthesis protein B | No |
| entD | D21A | 62 | C -> A | Substitution | 44 | 4'-phosphopantetheinyl transferase (enterobacin biosynthesis complex) | No |
| cysD | G291S | 871 | T -> C | Substitution | 41 | Sulfate adenylyltransferase subunit 2 | No |
| lsrK | T403S | 1,207 | A -> T | Substitution | 35 | Autoinducer 2 kinase LsrK | No |
| rscC | I362T | 1,085 | G -> A | Substitution | 38 | Sensor histidine kinase RcsC | No |
| udk | D74H | 220 | G -> C | Substitution | 62 | Uridine kinase | No |

Deletions of the following genes:
|  |  |  |
| --- | --- | --- |
| lipA | Lipoyl Synthase | Yes |
| cusF | Cation efflux system protein | No |
| cusB | Cation efflux system protein | No |
| cusA | Cation efflux system protein | No |
| pagP | Lipid A palmitoyltransferase | N.D. |
| cspE | Cold shock-like protein | No |
| crcB | Putative fluoride ion transporter | No |
| yafV | 2-oxoglutaramate amidase | N.D. |
| tatE | Sec-independent protein translocase protein | No |
| hypothetical protein | hypothetical protein | N.D. |

### Recombinant bacterial strains, cloning, and plasmid preparation

The ASKA library is a comprehensive GFP-tagged *E. coli* K-12 ORF plasmid library available from the National BioResource Project. The ASKA clone library is based on the E. coli K-12 strain AG1 and individual genes were cloned into the pCA24N vector (see Table 1) (26). The *E. coli* strain K-12 carrying pCA24N::*hemL, hemN, hemF, fes, fepC, araC, cysD, udk, pta, mprA, cusA, cusB, cusF, cspE, crcB, tatE, lipA, entD, tolC, rnr, hyfB* and pCA24N plasmid itself were obtained from the National BioResource Project. These *E. coli* strains were grown at 35° C for 18 h in TSY broth (Remel) with chloramphenicol (50µg/ml). Plasmids were isolated using the ZymoPure Plasmid MiniPrep kit (Zymo Research) and then transformed into the clinical strains of *E*.*coli, C*.*freundii*, and *E. hormaechei*.

One of the affected genes, *pqqB*, in the SCV *E. hormaechei* isolates was unavailable from the ASKA strain library as an ortholog of this gene is not present in *E. coli* K-12. This gene was amplified from the NCV *E. hormaechei* isolate with CloneAmp HiFi polymerase (Takara) and the following primers: 5’-TCC GGC CCT GAG GCC TAT GGC CTT TAT TAA AGT CCT CGG TTC C-3’ and 5’-TCC TTT ACT TGC GGC CGG GGT CCT GAA GCG TGA TGT TCA T- 3’. These PCR products were cloned into pCA24N with a C-terminal GFP tag using the In-Fusion HD enzyme kit (Takara). Clones were selected on TSA plates with 50µg/ml chloramphenicol. Sanger sequencing was conducted on the resulting plasmid to confirm cloning.

### Chemical and genetic rescue and cross-feeding

Unlike their NCV co-isolates, SCVs cannot grow on M9 minimal media. We took advantage of this to study the additional nutritional requirements of SCVs and determine which mutations were casual for the auxotrophic phenotypes. To study the nutritional requirements of each SCV, a 0.5 McFarland solution of the isolate was plated on M9 media. A disk impregnated with heme (Remel), δ-aminolevulinic acid (Oxoid), L-glutamate (Sigma-Aldrich, 30 mM), lipoic acid (Sigma-Aldrich, 5 µg/mL) or pyrroloquinoline quinone (Sigma-Aldrich, 3 µM) was placed onto the media. For each chemical rescue, the corresponding NCV co-isolate and a blank disk were included as controls. The M9 plates were examined after incubation for 20-24 hours at 35°C. Of note, NCV and SCV *C. freundii* carrying pCA24N were used in place of the untransformed NCV and SCV *C. freundii* due to the instability of the SCV isolate.

To examine the causal mutations responsible for the SCV phenotype, 0.5 McFarland solutions of each of the complemented strains were plated on to M9 media (Teknova). The M9 plates were examined after incubation for 20-24 hours at 35°C. Lastly, we examined whether the NCV could restore the growth of its SCV co-isolates by growing the strains adjacent to one another on M9 media. A 0.5 McFarland standard of each isolate was streaked on an M9 agar plate. This plate was examined after incubation for 20-24 hours at 35°C. Anaerobic cultures were also performed at 35°C using the AnaeroPack system (Mitsubishi Gas Chemical) and growth was observed at 48 hours.

### Preparation of competent cells and electroporation

To prepare electro-competent cells of *E*.*coli, C*.*freundii*, and *E. hormaechei*, 5 mL of TSY broth inoculated with a single colony was grown overnight with vigorous aeration (150 rpm/min) at 35° C. The following day, 30µl of overnight culture was diluted into 15ml of SOC media and grown at 37° C with constant shaking (180 rpm/min) until 0.5-0.8 OD600. Cells were harvested by centrifugation at 2000g for 10min at 4°C and washed twice with 10ml sterile ice-cold 10% glycerol. The supernatant was removed and the cell suspension was concentrated 50-fold in 3% glycerol.

For bacterial transformation, 100uL of electrocompetent cells were mixed with 100ng DNA in 0.1cm cuvettes. Electroporation was carried out using a Gene Pulser with the following parameters: 2.5 kV, 25 µF and 200 Ω for the NCV and SCV *E. coli*, NCV and SCV *E. hormaechei* and SCV *C. freundii*. The following parameters were used to transform the NCV *C. freundii*: 2.5 kV, 25 µF and 600 Ω. Immediately after pulsing, 0.9 ml of SOC media was added to each cuvette, the cell suspension was transferred to a test tube and then was incubated for 30min at 37°C with constant shaking. Transformed clones were selected on TSA plates with 50µg/ml chloramphenicol.

### Data availability

All assemblies are available from NCBI BioProject PRJNA523376 (https://www.ncbi.nlm.nih.gov/bioproject/?term=PRJNA523376).

## Results

### Clinical cases and isolates

Case histories are depicted in Figure 1 and described in the Materials and Methods. Briefly, the paired *E. coli* isolates were from a urine culture on a 6-week old otherwise healthy term male infant with fever. His white blood cell count was elevated at 21,600/ml and a 2+ urine leukocyte esterase at the time of emergency visit. The *C. freundii* isolates were from a urine culture on a 2-month old female infant with complex urological anomalies for duplicated collecting system and grade 3 vesicoureteral reflux. Her urine leukocyte esterase was 3+ with elevated red and white blood cells in urine at the time of culture. The *E. hormaechei* isolates were from multiple blood cultures, both arterial line and peripheral venous draw, spanning 3 days on a 6-year old male child with end stage renal disease receiving hemodialysis and multiple prior bloodstream infections in the prior year. Of note, these isolates were originally resulted out as *E. cloacae* based on MALDI-TOF species identification.

**Figure 1.**
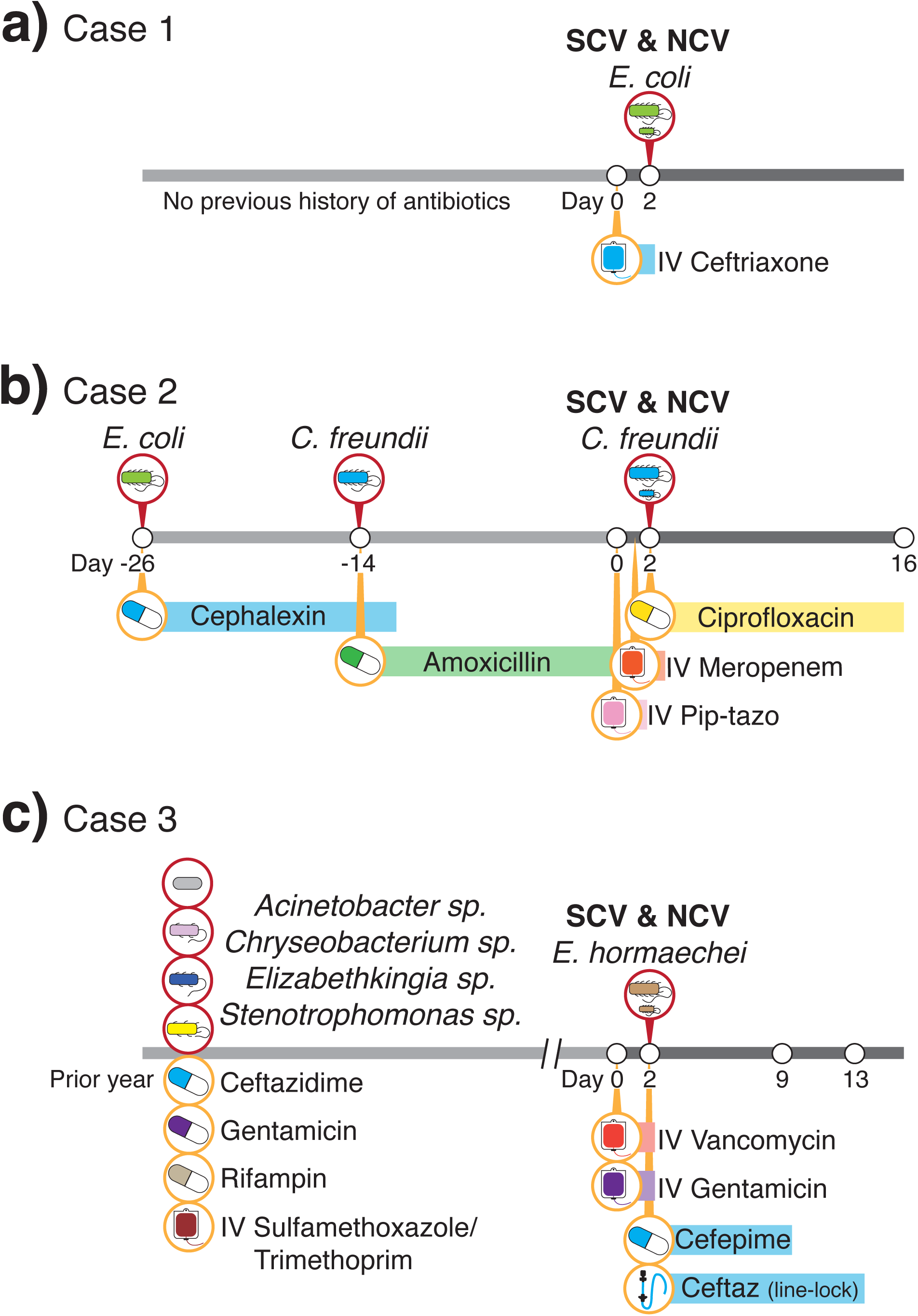
Case histories. Relevant past clinical microbiological and antibiotic selective pressures are indicated in the line histories for the isolation of NCVand SCV in Escherichia coli (a), Citrobacter freundii (b), and Enterobacter hormaechei (c).

### Antimicrobial resistance pattern is explained by ampC

Co-isolation of both NCV and SCV of the *Enterobacteriaceae* strains were common to all 3 cultures during routine culture workups. The antibiotic susceptibility pattern for the three NCV isolates is shown in Table 1 and followed expected patterns of resistance given the case histories. All three corresponding SCV isolates failed to grow on MH medium for susceptibility testing.

Genomic sequencing of the NCV isolates confirmed the presence of chromosomal *ampC* in all isolates, explaining the antibiotic susceptibility patterns recovered. Analysis of contigs with higher copy number revealed one small 4kb plasmid in *Escherichia coli*, three small 2.5 - 4kb plasmids in *Enterobacter hormachei*, as well as >80kb of phage sequence in the *Citrobacter freundii* assembly but no plasmids. No specific antimicrobial resistance genes were contained on these plasmids.

### Whole genome sequencing of paired isolates reveals parsimonious variants accounting for small colony phenotype

The *Escherichia coli* NCV assembly yielded 81 contigs >200 bp with an N50 of 360,223 bp. Mapping of the *Escherichia coli* SCV reads to the NCV assembly yielded only 4 variants, including 2 variants in heme-related genes (Table 2). The 272-amino acid enterobactin siderophore transport system ATP-binding protein (*fepC*) gene had an internal stop codon at amino acid 169. The *hemL* gene had a W59R coding change and the *hemF* gene had an in-frame 6-bp deletion resulting in the loss of an arginine and glutamic acid at amino acids 132 and 133. In addition, the transcriptional repressor *mprA* gene had a P84S coding change. No intergenic variants were recovered in the *Escherichia coli* SCV strain relative to the NCV strain.

Sequencing the *Citrobacter freundii* NCV strain yielded an assembly of 220 contigs >200 bp with an N50 of 61,653 bp. Mapping of the *Citrobacter freundii* SCV reads to the NCV assembly yielded a total of 6 variants (Table 2). Two of these variants were related to heme-producing genes. Most notable among these were a 1 bp deletion in the glutamate-1-semialdehyde 2,1-aminomutatase (*hemL*) gene that resulted in frame shift and stop codon at amino acid 75, as well as a premature stop codon at amino acid 918 of the enterochelin esterase (*fes*) gene. An additional heme-related variant included an intergenic mutation of unclear significance, 181 bp upstream of the aerobic coproprohphyrinogen-III oxidase (*hemF*) gene. The three remaining variants resulted in coding changes in genes unrelated to heme production, including a Q303L mutation in the phosphate acetyltransferase (*pta*) gene, a P778T mutation in the major subunit precursor of the nitrate-inducible formate dehydrogenase (*fdnG*) gene, and a P7L mutation in an arabinose operon regulatory protein (*araC*) gene.

The *Enterobacter hormaechei* NCV assembly yielded 60 contigs longer than 200 bp. Mapping of the *Enterobacter hormaechei* SCV reads to the NCV assembly yielded 7 single nucleotide coding variants, one of which was in a gene related to heme production and one involved in enterobactin synthesis. The oxygen-independent coproporphyrinogen-III oxidase (*hemN*) gene had a L366Q mutation, while the 4’-phosphopantetheinyl transferase (*entD)* involved in enterobactin synthesis complex had a D21A mutation. Multiple unrelated coding changes were found between the SCV and the NCV *Enterobacter hormaechei* strains, including a G291S mutation in the sulfate adenylyltransferase subunit 2 (*cysD*) gene, a C224W mutation in the coenzyme PQQ synthesis protein B (*pqqB*), a I362T mutation in the sensor histidine kinase (*rcsC*), a T403S mutation in the autoinducer 2 kinase (*lsrK*), and a D74H mutation in uridine kinase (*udk*). An intergenic G->A mutation 45 bp downstream of a hypothetical protein was also identified. A 10.2kb chromosomal deletion disrupting 11 genes was also found in the SCV assembly as compared to the NCV assembly. These genes included lipoyl synthase (*lipA*), cation efflux system locus (*cusA, cusB, cusF*), lipid A palmitoyltransferase (*pagP*), cold shock-like protein (cspE), putative fluoride ion transporter (*crcB*), Sec-independent protein translocase (*tatE*), 2-oxoglutaramate amidase (*yafV*), and two hypothetical proteins.

### Chemical and genetic rescue reveals causal SCV lesions

To determine which of the above lesions were responsible for the defects in growth observed *in vitro*, we performed chemical and genetic rescue experiments on the three SCV clones. We also tested the ability of the NCV isolates to rescue SCV growth by cross-feeding to further confirm that a diffusible factor was responsible for limited growth. All experiments were performed on M9 minimal medium.

*E. coli* SCV prototrophic growth was restored with overexpression of *hemL*, but not *hemF, fepC*, or *mprA* (Figure 2a). SCV growth was also rescued by the presence of heme (Figure 2b) or δ-aminolevulinic acid (ALA) (Figure S1a), but not L-glutamate, lipoic acid, or pyrroloquinolone quinone (PQQ) (Figures S1b-d). The SCV clone was also able to cross-feed from NCV (Figure 2c). These results are all consistent with the genetic deletion in *hemL* being responsible for the limited growth in our E. coli clone.

**Figure 2.**
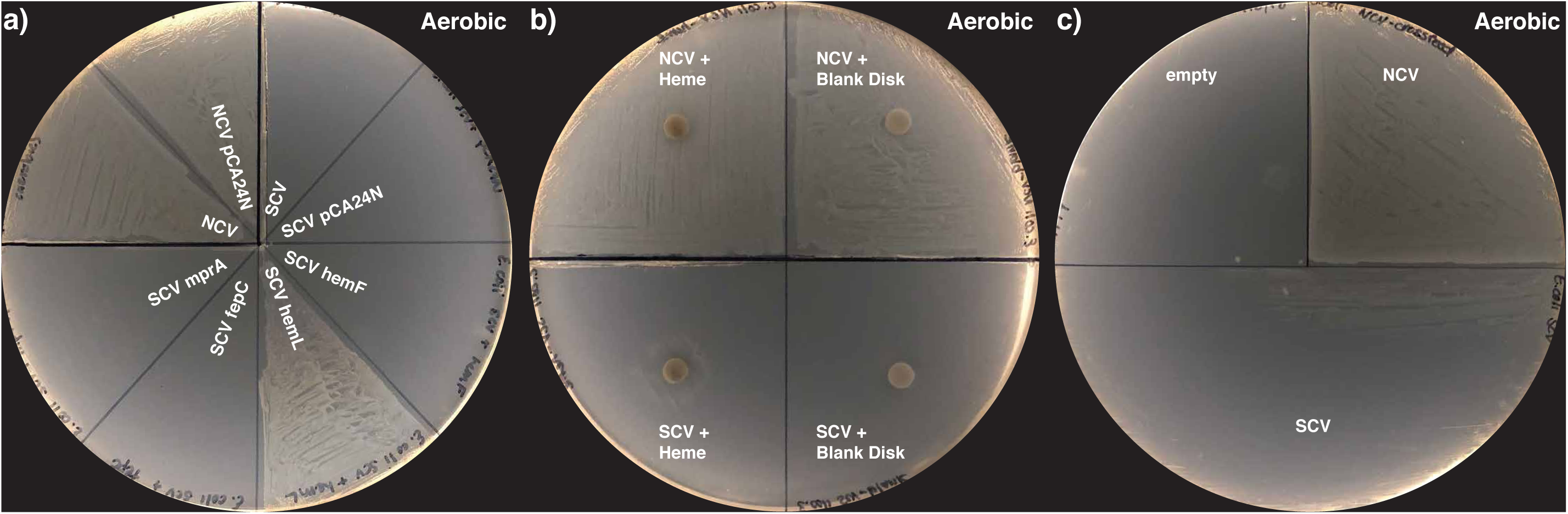
Heme biosynthesis lesion as cause of small colony phenotype in *Escherichia coli* isolated from urinary tract infection. Genomic sequencing of the paired NCV and SCV isolates revealed genomic lesions in *fepC, hemF, hemL*, and *mprA*. A) Only genetic rescue with *hemL* rescued normal growth from the *Escherichia coli* SCV. B) Chemical rescue with heme partially restored normal growth in *Escherichia coli* SCV. C) Cross-feeding from *Escherichia coli* NCV partially restores growth of SCV, consistent with a diffusible factor required for growth. All plates in this figure were incubated under aerobic conditions.

SCVs often revert to the NCV phenotype when serially passaged *in vitro*. We identified an *E. coli* SCV that reverted to a normal growth phenotype over the course of our study. We performed WGS on this SCV revertant to understand the individual mutations that resulted in wild type growth. This isolate had an NCV-like *hemL* sequence, but retained the SCV coding mutations in *hemF, fepC* and *mprA*. No additional mutations were observed in the reverted isolate. This further supports the results of our chemical and genetic rescues, and further highlights the importance of an intact *hemL* for normal growth.

Genetic and chemical rescue of *C. freundii* SCV growth showed similar results. *C. freundii* SCV prototrophic growth was restored with overexpression of *hemL* or *fes*, but not *araC* or *pta* (Figure 3a). The same chemical rescue results were seen as in the *E. coli* SCV, as heme (Figure 3b) or ALA (Figure S2a) were able to restore growth, but L-glutamate, lipoic acid, or PQQ failed to do so (Figure S2b-d). Moderate cross-feeding rescue with co-culture with the NCV clone was observed (Figure 2c). These results also indicate that deficiencies in heme synthesis and iron transport were responsible for the small colony growth phenotype seen in our *C. freundii* isolate.

**Figure 3.**
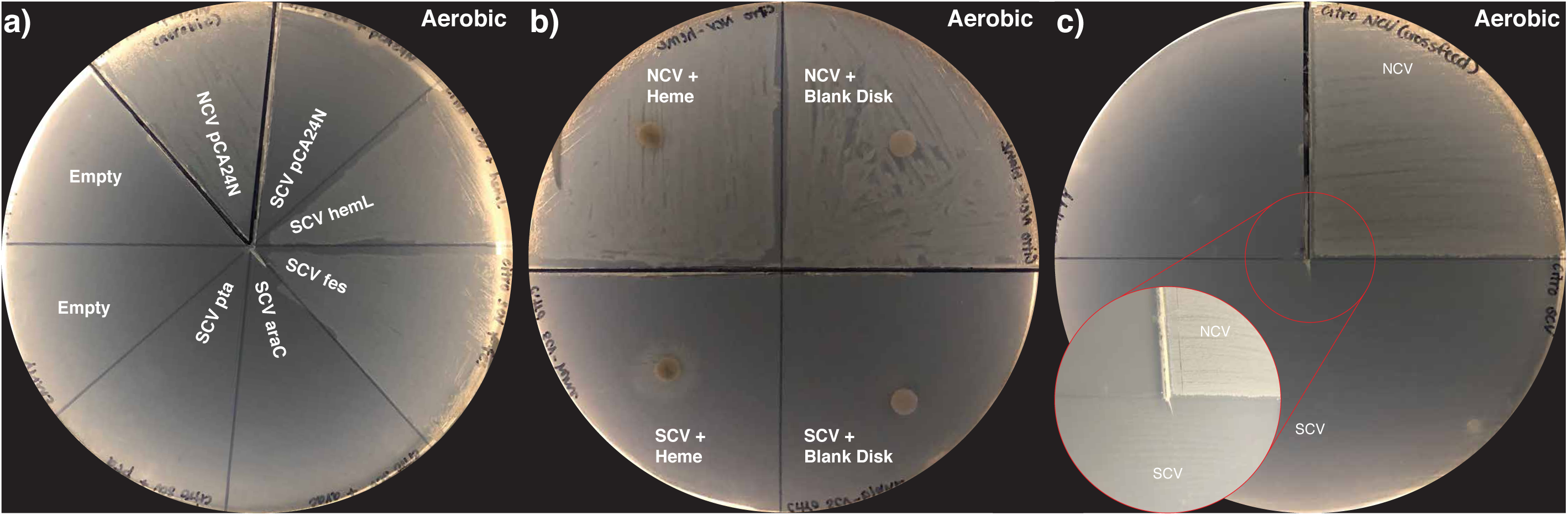
Heme biosynthesis along with iron availabiity lesion as cause of small colony phenotype in *Citrobacter freundii* isolated from urinary tract infection. Genomic sequencing of the paired NCV and SCV isolates revealed genomic lesions in *araC, fdnG, fes, hemL, pta*, and in the intergenic region upstream of the *hemF* gene. A) Genetic rescue with *fes* and *hemL* rescued normal growth from the *Citrobacter freundii* SCV. B) Chemical rescue with heme restored normal growth in *Citrobacter freundii* SCV.C) Cross-feeding from *Citrobacter freundii* NCV partially restores growth of SCV, consistent with a diffusible factor required for growth. All plates in this figure were incubated under aerobic conditions.

Over the course of the rescue experiments, the *C. freundii* SCV also reverted to normal growth. This clone had NCV-like *fes* and *hemL* sequence, while the same SCV coding mutations were seen in *araC, fdnG*, and *pta*, as well as the intergenic mutation upstream of *hemF*. The *C. freundii* revertant also had a new R100H mutation in the cytochrome bd-I ubiquinol oxidase subunit 2 gene (CydB) relative to both the SCV and NCV clones. These results further confirm the genetic and chemical rescue performed above, illustrating the importance of the heme pathway for normal growth.

Finally, despite also containing lesions in the heme biosynthesis pathway, the *E. hormaechei* SCV produced a radically different pattern of chemical and genetic rescue. Here, overexpression of *lipA* was the only gene that yielded prototrophic growth (Figure 4a), while all other disrupted genes failed to rescue growth (Figures 4a, S3a-c). Complementation with lipoic acid restored growth (Figure 4b) along with co-culture with *E. hormaechei* NCV (Figure 4c), while PQQ, heme and its biosynthetic intermediates L-glutamate and ALA failed to increase SCV growth (Figures S3d-g). These results conclusively demonstrate that disruption of the lipoylation pathway was responsible for the small growth phenotype in our *E. hormaechei* SCV isolate. We did not observe reversion to normal growth for the *E. hormaechei* SCV.

**Figure 4.**
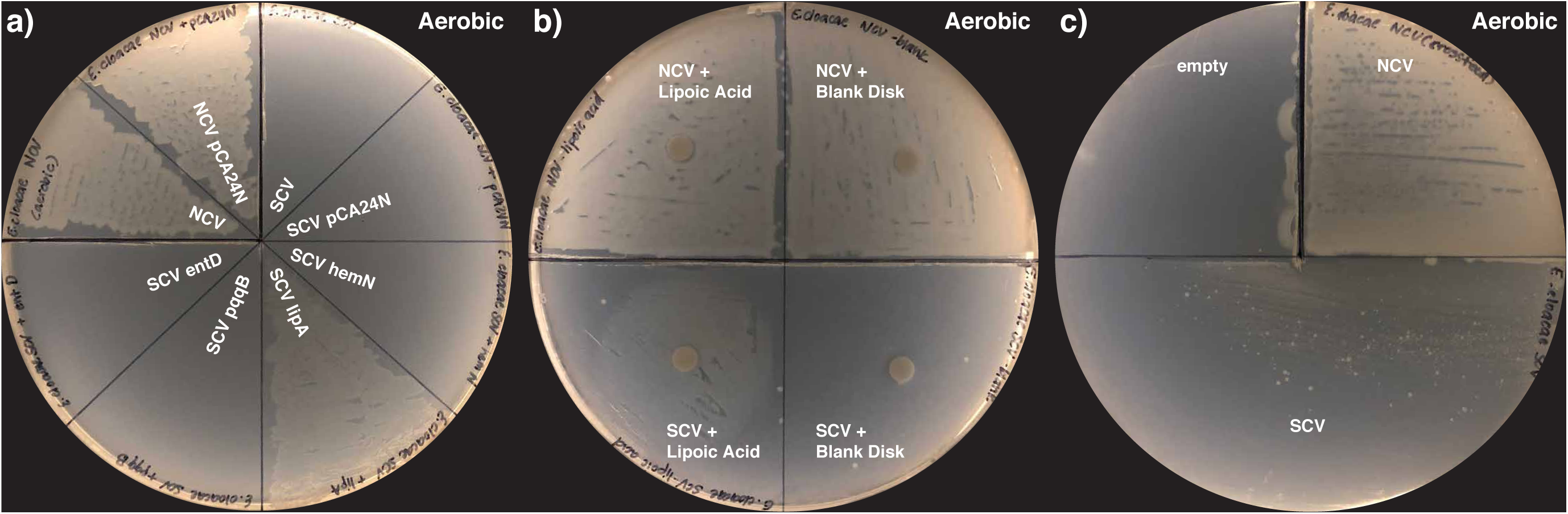
Lipoic acid biosynthesis as cause of small colony phenotype in *Enterobacter hormaechei* isolated from a bloodstream infection in a patient with end-stage renal disease. Genomic sequencing of the paired NCV and SCV isolates revealed multiple genomic lesions including single nucleotide substitutions in *entD, hemN*, and *pqqB* along with large-scale rearrangements leading to disruption of the *lipA* gene. A) Genetic rescue with *lipA* restored normal growth from the *Enterobacter hormaechei* SCV. B) Chemical rescue with lipoic acid restored normal growth in *Enterobacter hormaechei* SCV. C) Cross-feeding from *Enterobacter hormaechei* NCV restores growth of SCV, consistent with a diffusible factor required for growth. All plates in this figure were incubated under aerobic conditions.

### SCV demonstrate prototrophic growth under anaerobic conditions

Based on recurrent isolation of mutants in aerobic respiration pathways, along with the genetic and chemical rescue experiments demonstrating their causality, we hypothesized that SCV isolates might not demonstrate growth defects under anaerobic conditions. We cultured NCV and SCV isolates for each of the three species along with the transformed genetic rescue clones under anaerobic conditions for 48 hours. Anaerobic conditions rescued SCV growth in all cases (Figure 5a-c, S4h-j). These results were also independent of every genetic construct transformed.

**Figure 5.**
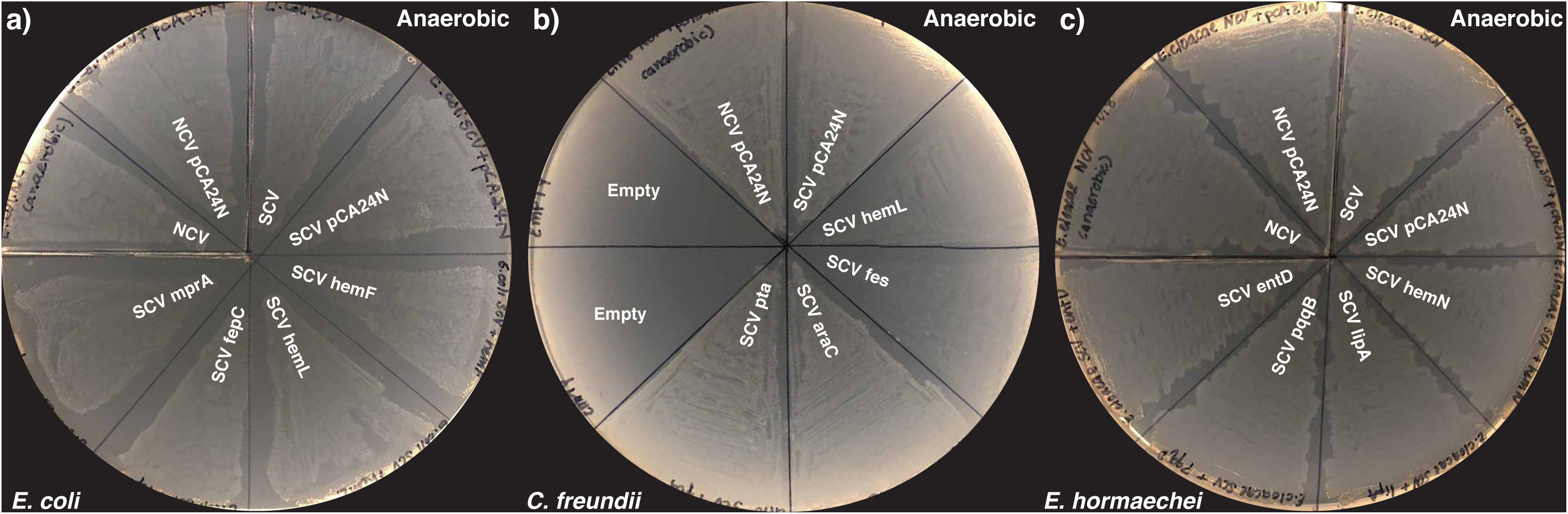
Anaerobic growth rescues growth for small colony variants in *E. coli* (A), *C. freundii* (B), and *E. hormaechei* (C) isolates.

## Discussion

Here, we used genomic screening of paired isolates to understand the molecular mechanism of the small colony growth phenotype in Gram-negative bacteria encountered in the clinical microbiology laboratory as well as bacterial persistence *in vivo*. Unlike the NCVs, all three corresponding bacterial SCV isolates were auxotrophic, thus unable to grow in glucose only M9 medium. Using genetic, chemical rescue, as well as NCV cross-feeding, we found heme-production pathway lesions to be responsible for the SCV phenotype in two of the three isolates, while lipoic acid synthesis was responsible for the third. In the *E. coli* SCV, both W59R in *hemL* and two-amino acid deletion in *hemF* could impact heme production, but only complementation of hemL rescued growth on M9 media. In the SCV *C. freundii*, the truncation of *hemL* blocked ALA production. In *E. hormaechei* SCV, the *lipA* gene was interrupted by large-scale genomic deletion and, correspondingly, complementation with *lipA* or lipoic acid restored prototrophic growth. Intriguingly, growth of the SCVs was not impaired under anaerobic conditions, consistent with the role of heme as an essential cofactor for the electron transport chain and lipoic acid’s role in the Krebs cycle. Of note, this is the first report of whole genome comparison of paired NCV and SCV *Enterobacteriaceae* isolates associated with clinical bloodstream and urinary tract infections from pediatric patients and the first report of detection of gene lesions associated with bacterial iron acquisition. Gene truncations in enterochelin esterase (*fes*) in the SCV *C. freundii*, a putative siderophore transport system ATP-binding protein (*fepC*) in the SCV *E. coli*, and D21A in *entD* of *E. hormaechei* were novel findings with potential implications for bacterial lifestyle changes upon host selection. Notably, despite the significant genetic lesions recovered here – two truncations and a coding mutation never previously recovered in any *Enterobacter hormaechei* -- complementation with *fepC* and *entD* genes from *E. coli* failed to rescue prototrophic growth. Future work will need to characterize the effect of these mutations on protein function, as it is possible that these mutations result in dominant negative phenotypes.

The antibiotic exposure history or the choices for certain specific agent(s) used for the treatment in the 3 patients at the time could not explain a common pattern of selective pressure for the emergence of bacterial SCV. The patient underlying illnesses also ranged from the first episode of a potential *E. coli* urosepsis in a newborn, a recurrent *C. freundii* UTI in a 2-month old, and a presumed *E. hornaechei* occult renal system infection in a 6-year old patient with end stage renal disease pending kidney transplant. The second and the third cases both had significant prior antimicrobial exposure to multiple classes of agents with beta-lactams being the most common agent, but first new born case had no prior treatment ever. Although prior exposure to aminoglycosides or sulfamethoxazole-trimethoprim has been reported as selective pressure for selection of bacterial SCVs (4, 27), these two agents were only used in the third case. Therefore, antibiotic use alone may not be the major contributor for SCV selection.

Pairwise sequencing of the NCV and SCV genomes for SCV mutational characterization was based on the assumption that both NCV and SCV descended from the same strain. While the NCV is likely the closest representative of the parental strain, SCV diverged from this strain with distinctive mutations that were absent in NCV. The functional M9 growth studies *in vitro* have clearly demonstrated that deficiencies in bacterial synthesis of heme, lipoic acid, and/or iron acquisition apparatus were the primary contributors to SCV auxotrophism. Our analysis suggests that both the oxidative respiration and iron acquisition may be counter-selected by the host during the subacute or perhaps chronic infections. Moreover, the extent of SCV genomic mutations could vary depending on the clinical course of the infective illnesses. For example, the *E. coli* SCV urinary tract isolate from a 6 week old infant had 4 SNPs, the repeat *C. freundii* SCV urinary tract isolate from a 2 month old patient with urological anomalies had 6 SNPs, and the blood stream *E. hormaechei* SCV isolate from a 6 year old patient with end stage renal disease had 8 SNPs plus an 11-gene deletion.

The selective loss of oxidative respiration and iron acquisition we observed in the SCV isolates sharply contrasts with conventionally held beliefs about microbial pathogens. Rather than rapidly dividing and completing fiercely for iron, SCVs take a unique approach for evading the host defense response, which includes iron starvation and oxidative stress (1, 2). In response to the iron-limiting condition, it has been well documented that the bacterial ferric uptake regulator (Fur) system is activated to strengthen microbial ability to capture iron for energetic growth and pathogenesis (3). Regardless of iron availability, *Enterobacteriaceae* are facultative anaerobes, and they are fully capable of growth under various oxygen tensions with oxidative respiration being significantly more robust for energetic growth *in vitro* (28). Therefore, change in oxygen tension itself in the infected host environment may not be the major selective pressure for SCV development. Our finding of mutations in oxidative respiration related genes was consistent with the overwhelming reports of bacterial SCV deficiencies in heme, menaquinone, and lipoic acid synthesis (4, 21, 29, 30). Thus, an overarching impression of the SCV degenerative mutations is their association with the essential elements of aerobic living which demands more iron (Figure 6) (31). The SCV isolates in this study, regardless of heme, lipoic acid, and/or iron transport deficiencies, were all able to grow anaerobically on M9 without nutrient supplementation. In fact, the selective loss of aerobic respiration under iron limiting and oxidative stress conditions is indicative of a microbial survival mechanism by “retreating to anaerobic habitats” in order to persist inside a single affected host without clonal dissemination into the host population (4, 32–34). Hence, resorting to anaerobic living may be an alternative response to iron limitation and oxidative stress (35). This alternative survival mechanism would be opposite to the well-known “superbugs” survival strategy of head-on iron competition and host population dissemination (36, 37). It also shares similarities with the immunoevasion strategies of tumors via hypoxia-induced immune exhaustion and T-cell anergy (38).

Iron is an essential nutrient for all forms of life (39). However, some free-living and obligate parasitic bacteria (e.g. *Borrelia burgdorferi*) employ a unique iron-independent redox-active metal such as manganese (Mn) to deal with oxidative stress, which allows them to bypass host iron defense (40). *E. coli* maintains a Mn-SOD (superoxide dismutase) system which is also regulated by Fur (41). All three SCV isolates contained an intact *sodA*, though expression and functional anti-oxidant activities were not examined. It is conceivable that the inactivation of iron uptake apparatus in bacterial SCVs may be indicative of bacterial transition into a parasitic lifestyle by means of iron bypass (40). Alternatively, the genetic lesions described here, particularly in *E. hormaechei*, could also be indicative of evolution to a more symbiotic lifestyle, as evidenced by the cross-feeding and gene decay (42). The co-obligate symbiotic bacterium *Serratia symbiotica*, known for its minimal genome (~143kb), has lost the ability to synthesize both heme and enterobactin and, additionally, lacks an iron uptake apparatus, similar to the changes observed in our SCV isolates (43, 44).

This study is limited by our focus only on mutations that were associated with auxotrophism on M9 medium without analyzing other mutations that could potentially affect many other aspects of bacterial functions. For example, a ~10.2-kb deleted fragment in *E. hormaechei* SCV contained *cusF, cusB*, and *cusA* in tandem where CusCFBA is a membrane-bound proton antiporter for Cu/Ag efflux (45). As this efflux system carries out ATP-demanding activities, we speculate that this function encounters the same counter-selective pressure as a result of reduced ATP production assuming SCV anaerobic lifestyle. The exact host factor(s) that was critical for selection of SCV mutations is still unknown. Ideally, host selective factors could be identified if we could recreate the bacterial SCVs from the wildtype using host materials for bacterial growth *in vitro* or in animal models.

Antimicrobial resistance and iron competition as mechanisms for bacterial survival have attracted a great deal of interest (46). These isolates are now readily recognized in the clinical microbiology laboratory due to their robust growth and detectable resistance phenotypes using standard *in vitro* testing. Clinical isolates such as SCVs that lack these growth characteristics may be easily missed due to their small colony size and failure to grow on standard susceptibility testing media. The role of these alternatively pathways of survival in clinical infections may be significantly underestimated (30). Future work must address the totality of bacterial survival mechanisms, including mechanisms of bacteria persistence via resistance to host iron immunity and oxidative stress by means of anaerobiosis and iron bypass.

## Acknowledgements

We thank National BioResource Project NBRP *E*.*coli* Strain, Japan for the *E. coli* plasmid library (https://shigen.nig.ac.jp/ecoli/strain/). We also thank Seattle Children’s Microbiology team for their effort in standardization of bacterial SCV identification and quality documentation of these paired isolates.

---

**Figure S1.**
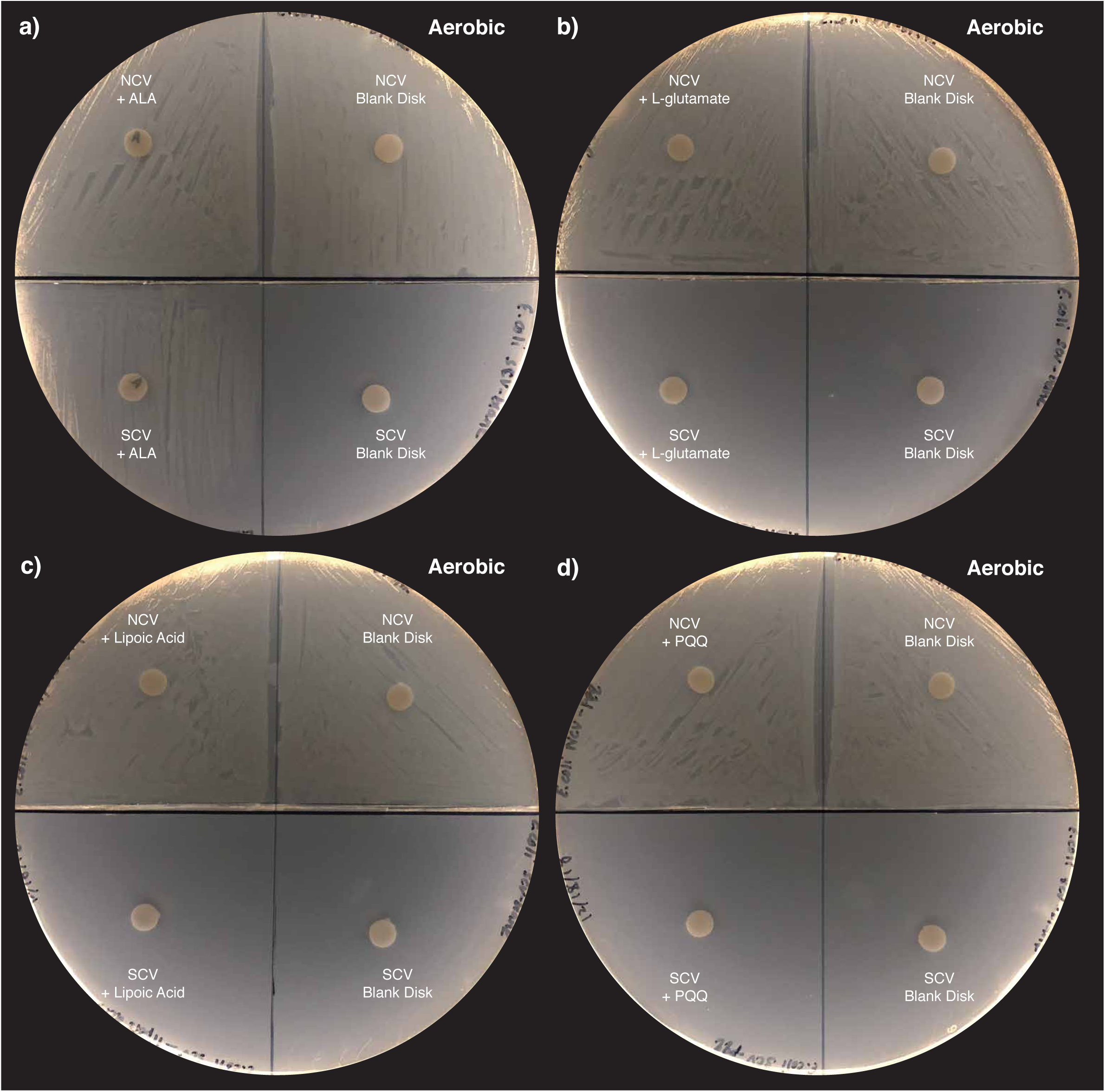
Chemical rescue of Escherichia coli SCV growth was successful with δ-aminolevulinic acid (a), but not with L-glutamate (b), lipoic acid (c), or pyrroloquinoline quinone (d).

**Figure S2.**
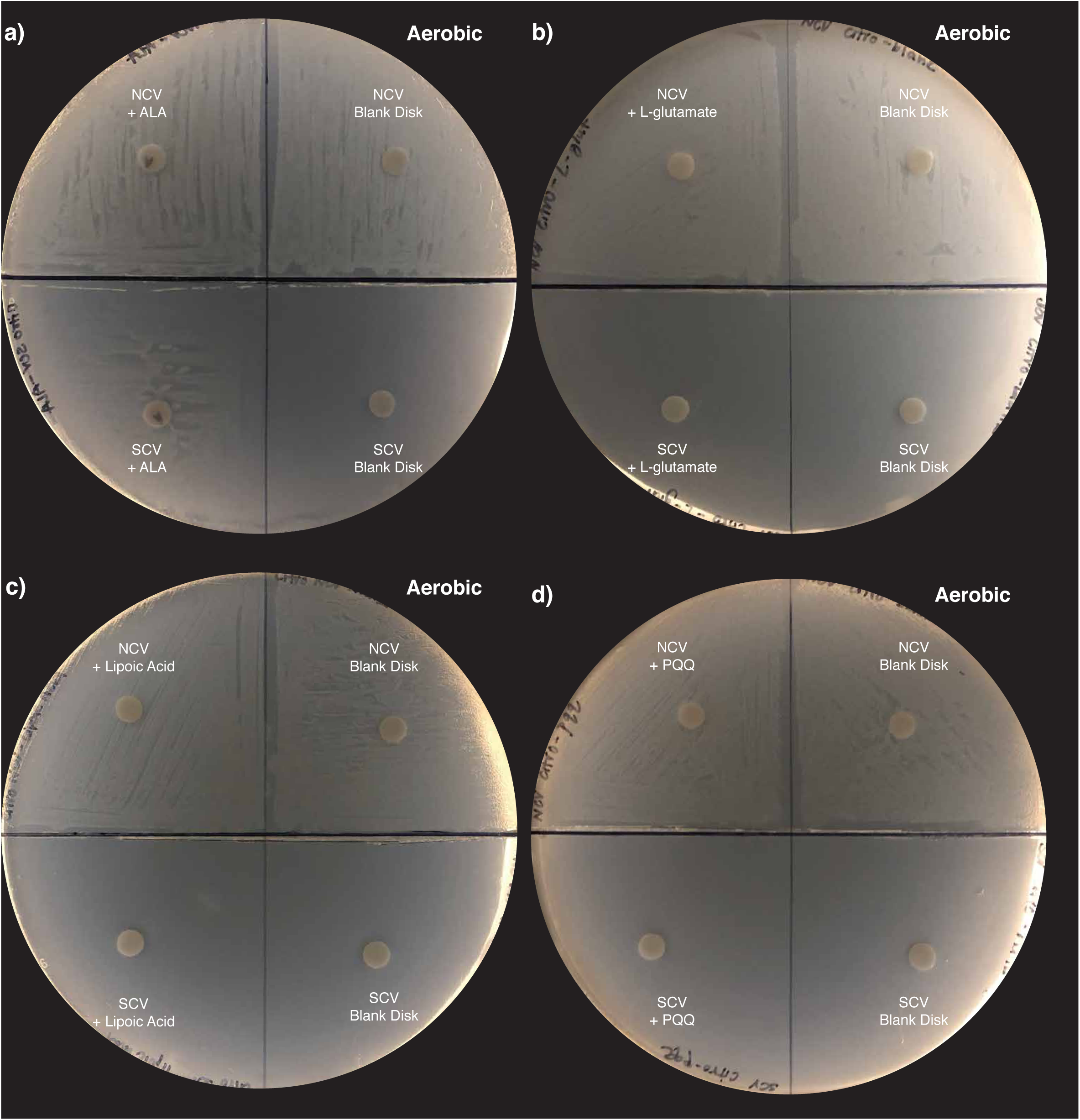
Chemical rescue of Citrobacter freundii SCV growth was successful with δ-aminolevulinic acid (a), but not with L-glutamate (b), lipoic acid (c), or pyrroloquinoline quinone (d).

**Figure S3.**
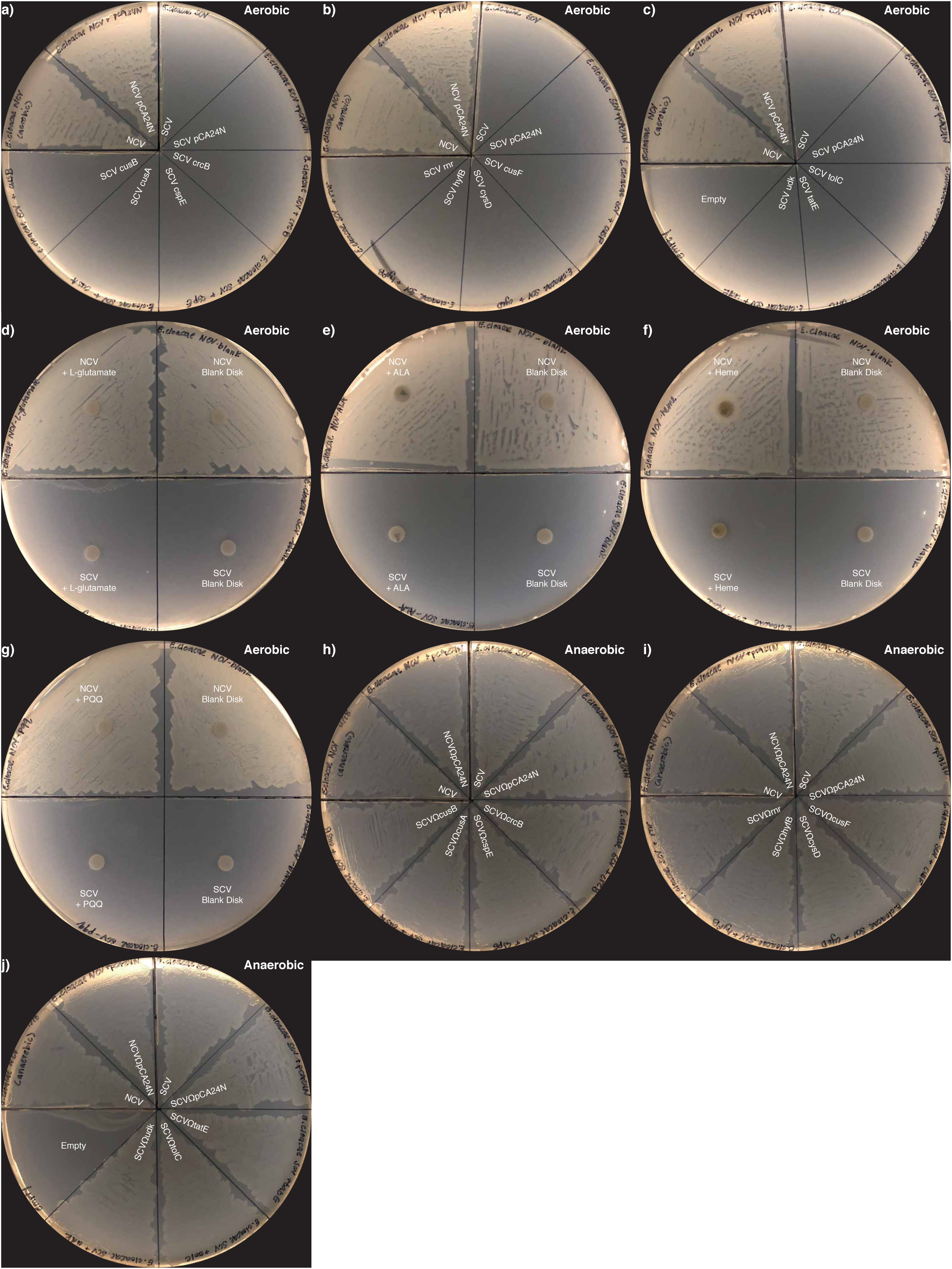
Failure to rescue Enterobacter hormaechei SCV growth with deleted and mutated genes crcB, cspE, cusA, cusB, cusF, cysD, hyfB, rnr, tatE, tolC, and udk under aerobic conditions (a-c). L-glutamate (d), δ-aminolevulinic acid (e), heme (f), and pyrroloquinoline quinone (g) failed to rescue SCV growth under aerobic conditions. SCV grew similar to NCV under all anaerobic growth conditions for each of the genetic rescues attempted (h-j).

